# annoSnake: a Snakemake workflow for taxonomic and functional annotation of metagenomes and metagenome-assembled genomes (MAGs)

**DOI:** 10.1101/2025.11.03.686227

**Authors:** Bastian Heimburger, Rebecca Clement, Tamara R. Hartke

## Abstract

Omic technologies revolutionised research in microbiomics and helped decipher the complex community structure of gut microbiota, each comprising hundreds of species of bacteria, archaea, and protists. Even as sequencing costs decrease, understanding these communities remains a challenge. Here, we introduce annoSnake, a Snakemake workflow that facilitates taxonomic and functional annotation of metagenomes and metagenomic-assembled genomes (MAGs). This user-friendly and highly flexible workflow integrates state-of-the-art software packages, taking raw sequencing reads, assembling them into contigs, assigning taxonomies and annotating them using various popular protein databases. The workflow handles paired-end as well as interleaved read data, and provides publication-ready figures and tables for downstream analysis. annoSnake, as all Snakemake workflows, is highly scalable, reproducible, and portable, enabling execution across diverse high-performance computing environments with remarkable efficiency.

## Introduction

Metagenomics is increasingly popular due to cost-efficient high-throughput sequencing technologies (Lema et al. 2023, Nam et al. 2023). However, data analysis remains tedious and time-consuming (New and Brito 2020, Setubal 2021, Zhou et al. 2022). Typically, standalone programs such as MEGAHIT (Li et al. 2015), Prokka (Seeman 2014) and Diamond (Buchfink et al. 2021) are used to perform a single task (e.g. genome assembly, taxonomic or functional annotation) en route to fully annotated metagenomic contigs, making researchers navigate multiple data formats and database setups. Although some (semi)automatic pipelines exist (Kultima et al. 2016, Dong and Strous 2018), no existing pipeline covers assembly and annotation of metagenomes and MAGs simultaneously.

Here, we introduce annoSnake, a Snakemake workflow that streamlines taxonomic and functional annotation of metagenomes and MAGs, from raw sequencing reads to fully annotated metagenomes and MAGs. Users select input read format (paired-end or interleaved) and annotation databases (eg. Pfam, KEGG, CAZy), which are automatically downloaded and set up. Resulting CSV tables and figures (PDF and HTML formats) are ready for downstream analysis and publication.

Snakemake workflows, like annoSnake, ensure reproducibility (dependencies are explicitly defined), scalability (designed to run on cluster environments), and testability (debugging is easy: single tasks are encapsulated into separate rules), allowing researchers to concentrate on interpreting results rather than bioinformatic problems. We demonstrate annoSnake here with a termite gut metagenomics project.

### Description

To start, users install Snakemake (Mölder et al. 2021; https://snakemake.readthedocs.io/en/stable/getting_started/installation.html), clone the annoSnake GitHub repository, and configure the .*/profile/params*.*yaml* file (see below). As the workflow is optimised for High Performance Computing (HPC) clusters using the Slurm workload manager (Yoo et al. 2003), users modify the .*/profile/config*.*yaml* file according to their cluster environment specifications.

Users provide either paired-end or interleaved sequencing data (in gzipped format), conforming to the naming scheme (i) $SAMPLE_R1/_R2.fastq.gz or (ii) $SAMPLE.fastq.gz. There is no need to trim or filter reads in advance. First, annoSnake creates all necessary Conda environments and sets up protein databases (i.e., download of databases and configuration, if necessary). Then, rules are processed consecutively to produce desired output files from input files (Fig. 1).

**Figure 1.**
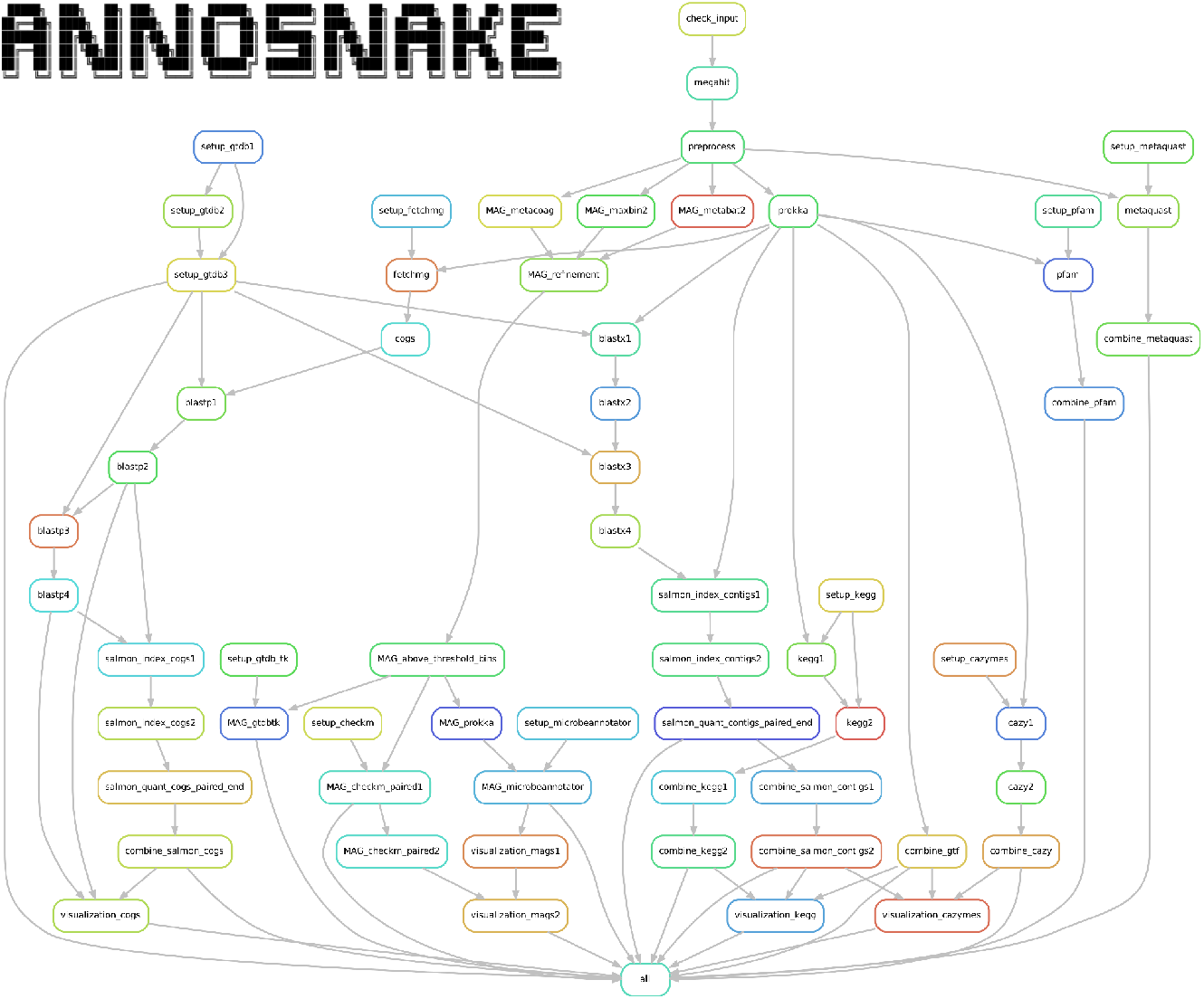
Full directed acyclic graph of jobs for an annoSnake run using paired-end sequencing reads. The workflow includes download and setup of databases; metagenome and MAG assembly; taxonomic, structural (protein-coding sequences, tRNAs, mRNAs), and functional (KEGG, CAZymes, Pfam) annotation of metagenomes and MAGs; and creation of tables and figures.

In the worked example, annoSnake detects enzymes associated with lignocellulose digestion. However, annoSnake can facilitate searches for any KEGG pathway (e.g. digestive enzymes, immune response proteins). Users simply adjust the files (i) *keggid_to_gene_name*.*csv* and (ii) *keggid_to_genes_pathway*.*csv* within ./workflow/rules/scripts/ to include relevant KEGG IDs and gene names to query.

### Configuration files

.*/profile/params*.*yaml* is the main configuration file. Users specify input and output directories and library type (‘paired-end’ or ‘interleaved’). Other options include (i) MAG assembly, (ii) databases for functional protein annotation, and (iii) corresponding cut-off E-values.

.*/profile/config*.*yaml* handles Slurm job submission. Users specify the partition (or queue) on which jobs are executed. Note: for large data sets, it may be necessary to adjust the amount of memory per node and maximum wall-clock time.

### annoSnake workflow: step by step

#### Metagenome assembly

Raw reads are assembled with MEGAHIT v1.2.9 (Li et al. 2015), optimised for metagenome assemblies. Users specify in .*/profile/params*.*yaml* the minimum length of contigs (default: 1500 bp) to retain for downstream analyses. To account for the complexity of metagenomes, ‘--presets meta-sensitive’ is used as default. Fasta headers of assembled metagenome contigs are modified prior to downstream analyses to include the $SAMPLE name. Quality of metagenome assemblies is assessed with metaQuast (Mikheenko et al. 2016).

#### Taxonomic annotation

Taxonomic assignment of metagenome contigs is performed as follows: (a) Prokka 1.14.6 (in ‘--metagenome’ mode; Seeman 2014) identifies protein-coding sequences (CDS), rRNAs, and tRNAs used later in the workflow (see Fig. 1). (b) fetchMG v.1.2 (https://github.com/motu-tool/fetchMGs) extracts 40 single-copy marker genes present in most living organisms (Sunagawa et al. 2013). (c) Marker gene protein sequences are taxonomically assigned with DIAMOND (Buchfink et al. 2021) in ‘blastp’ mode. (d) Other predicted protein-coding sequences (in nucleotide format) are taxonomically assigned with DIAMOND in ‘blastx’ mode. Both annotations use GTDB database ver 202 (Parks et al. 2022) as default reference.

To retrieve taxonomic classifications of predicted protein-coding sequences, annoSnake implements a custom R script ‘gtdb_diamondlca.R’ (Arora et al. 2022).

#### Functional annotation

Metagenomic contigs assigned as bacteria or archaea by ‘blastx’ are annotated. Users choose the databases for functional annotation: (a) default for CDS with carbohydrate metabolising properties is Hidden Markov models (HMM) of CAZy domains deposited in the dbCAN database release 11 (Yin et al. 2012). (b) For hydrogenases, HMM searches against the Pfam database version 35 (Mistry et al. 2020). Results can be used to further classify catalytic subunits with the HydDB webtool (Søndergaard et al. 2016; not implemented in annoSnake). (c) KofamScan v1.3.0 (https://github.com/takaram/kofam_scan) reconstructs prokaryotic metabolic pathways (i.e. genes related to methanogenesis, reductive against the KEGG database; Kanehisa and Goto 2000, Kanehisa et al. 2023). Cut-off E-values (minimum significant hit) are specified in .*/profile/params*.*yaml*.

#### Abundance of gene families

Abundance of CDSs is quantified with Salmon v1.10.2 (Patro et al. 2017) using specific settings for metagenomic analysis which align raw sequencing reads to assembled contigs annotated as bacteria or archaea. Salmon adjusts for biases such as GC-content and gene length, producing CDS abundances as Transcripts per Million (TPM; Patro et al. 2017). In addition, Salmon quantifies relative abundance of bacterial and archaeal lineages using the 40 single-copy marker genes (see section ‘Taxonomic annotation’).

TPM values >1 (to exclude potentially unreliable data) are log-transformed for visualisation. Normalisation of TPM counts is performed *via* centered log-ratio (clr) transformation (Calle et al. 2019) to account for the compositional nature of metagenomic data and allow comparisons between genes and bacterial species (Xia and Sun 2021). The transformation is executed using R package propr (Quinn et al. 2017) with a pseudo count of 1 to handle zero values appropriately.

#### Metagenome-assembled genomes (MAGs)

Metagenome contigs are binned into MAGs with three different binning algorithms (in default mode): (i) MetaBAT version 2.10.2 (Kang et al. 2019); (ii) MetaCoAG v1.1.1 (Mallawaarachchi and Lin 2022); and (iii) MaxBin 2.2.7 (Wu et al. 2016). To increase contiguity and completeness of bins, annoSnake implements metaWRAP’s (v1.3; Uritskiy et al. 2018) ‘bin_refinement’ module to produce a consolidated bin set from the three algorithms. Users specify in ./profile/params.yaml minimum completeness and maximum contamination of MAGs to use for downstream analyses. Defaults are ≥50% completeness with ≤10% contamination, following Bowers et al. (2017).

MAGs passing quality control (CheckM 1.2.2; Parks et al. 2015) are taxonomically classified with GTDB-Tk v2.3.2 (Chaumeil et al. 2022) using GTDB database ver 214 as a reference (Parks et al. 2020). Gene prediction is performed by Prokka (Seemann 2014), using the ‘--metagenome’ option, accounting for highly fragmented metagenomes. Predicted protein sequences are annotated with MicrobeAnnotator (Ruiz-Perez et al. 2021) using DIAMOND sequence aligner and KofamScan. Pathway completeness is assessed based on presence/absence.

#### Databases

Databases are automatically downloaded and set up. Users can choose to use their own protein databases, which must be stored in the correct format. All databases consume approx. 100 Gb of memory.

#### Output

annoSnake outputs tables (in CSV format) and figures (PDF and HTML formats), depending on protein annotations and whether MAGs are assembled or not. Tables and figures are ready for downstream (statistical) analysis and publication. Figures are visualised with ggplot2 3.4.3 (Wickham 2016) and plotly 4.10.4 (Sievert, 2020) in R version 4.2.0 (R Core Team 2024).

### Empirical data

We highlight the functionality of annoSnake using low-coverage WGS data from the Australian *Amitermes* group (AAG). These derived termites exhibit diverse feeding habits and nesting behaviour (Krishna et al. 2013). Although major decomposers of dead plant material in Australia (Coventry et al. 1988, Noble et al. 2009, Evans et al. 2011), their microbiomes are virtually unknown (but see Platell 2019). The worked example uses paired-end Illumina NextSeq data from 31 colonies representing nine *Amitermes* and *Drepanotermes* species (for details see Tab. S1 and Supplementary Information); the output of the worked example can be found under .

## Results and Discussion

Although low-coverage WGS provided a limited number of reads (∼212x10^6^ total, average ∼7x10^6^ reads/sample), annoSnake efficiently identified major bacterial lineages (Fig. 2) and a large of set of metabolic-pathway genes (KEGG) and CAZYmes (Fig. 3 and 4) related to lignocellulose digestion. For this small dataset, we estimate annoSnake saved tens of hours of bioinformatics work compared to the same tools in a non-automated workflow. It started downstream analyses without human intervention as the necessary data passed quality controls, reducing server time while avoiding errors and frustration associated with manually wrangling files and changing formats.

**Figure 2.**
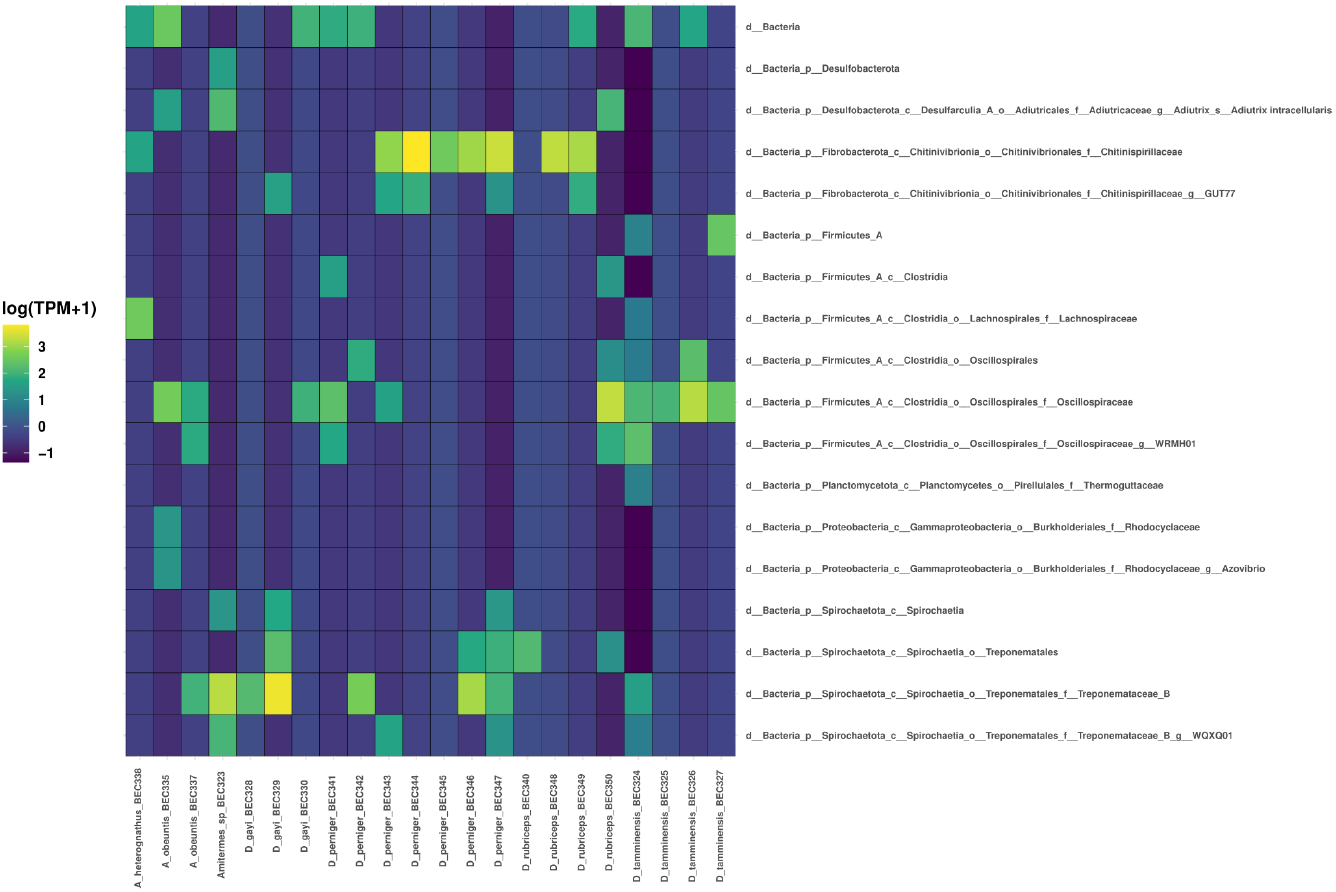
annoSnake output of relative abundance of bacterial lineages in termite gut metagenomes. Colour scale represents log transcripts per million (TPM), with lighter colours indicating higher relative abundances and darker colours indicating lower relative abundances.

**Figure 3.**
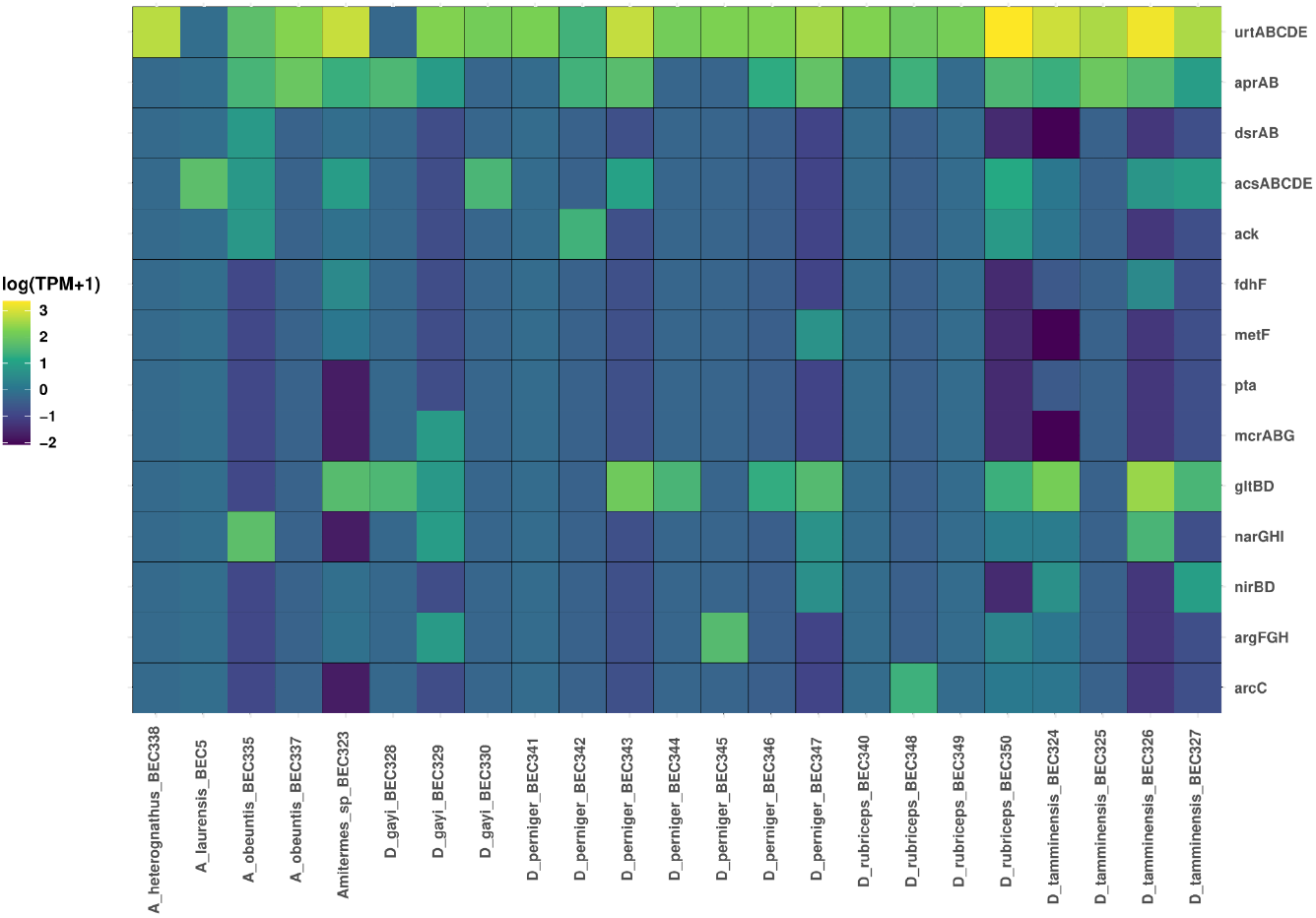
annoSnake output for relative abundance of metabolic pathway genes (KEGG) involved in lignocellulose digestion. Colour scale represents log transcripts per million (TPM), with lighter colours indicating higher relative abundances and darker colours indicating lower relative abundances.

**Figure 4.**
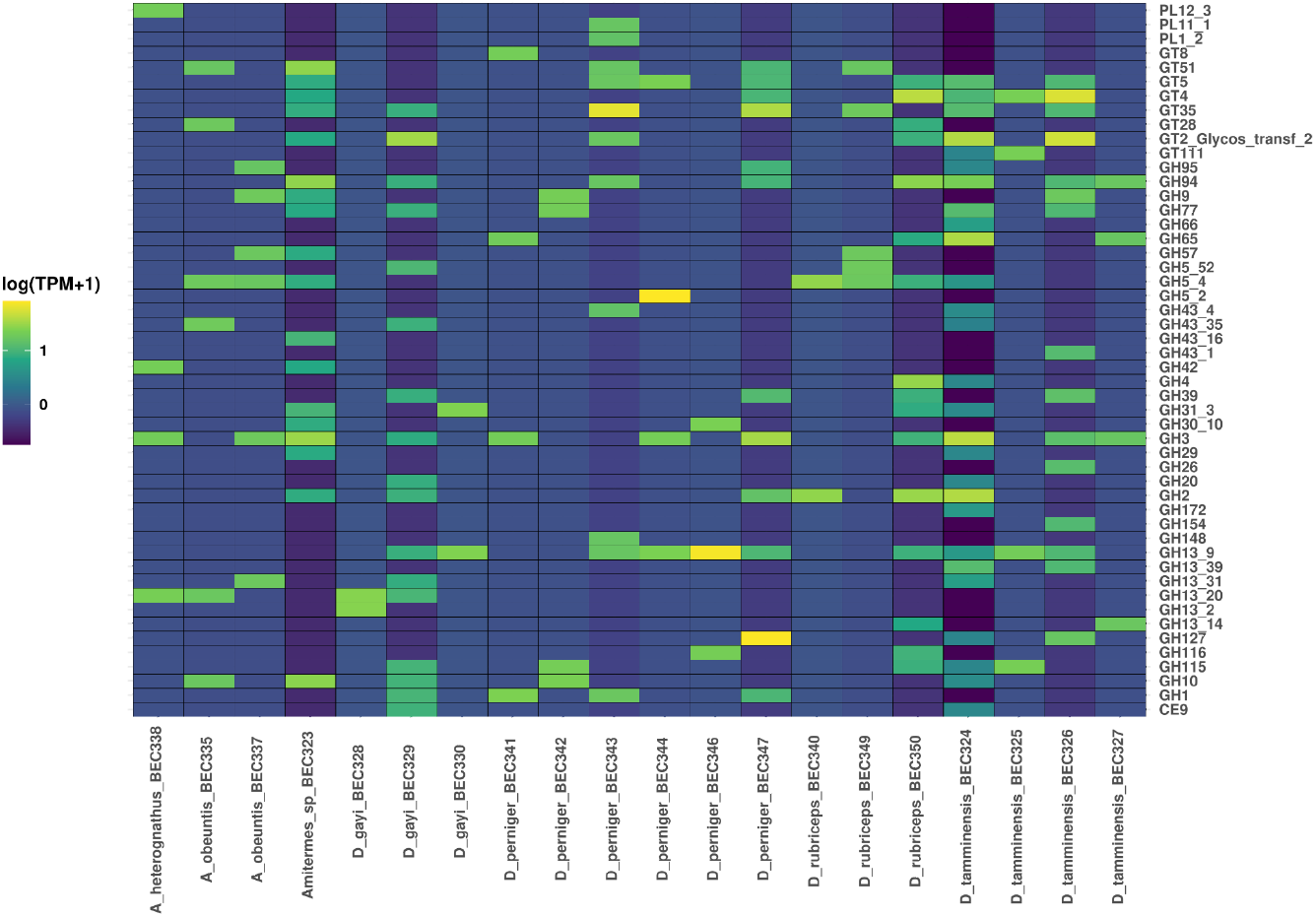
annoSnake representation of relative abundance of 50 most abundant CAZymes in termite guts. CAZYme annotation was performed by HMM searches against the dbCAN database release 11. Colour scale represents log transcripts per million (TPM), with lighter colours indicating higher relative abundances and darker colours indicating lower relative abundances.

Typically, grass- and wood-feeding termites have spirochete-dominated gut communities, while humus- and soil-feeders are rich in clostridia (Ottesen and Leadbetter 2011, Mikaelyan et al. 2015, 2016). annoSnake taxonomic annotation (Fig. 2) indicated that spirochetes indeed dominate most gut communities of the leaf litter and grass harvesting *Drepanotermes* (Watson and Perry 1981). Surprisingly, gut communities of *D. tamminensis* are heavily dominated by clostridia, with virtually no spirochetes or Fibrobacterota, as were communities of some *D. gayi* colonies. In the case of *A. heterognathus*, workers may have fed on dung in addition to their usual wood (Gay 1967, Ferrar and Watson 1970), resulting in a clostridia-rich community similar to facultative dung-feeder *A. wheeleri* (He et al. 2013). Unlike Arora et al. (2022), no archaeal lineages were found in any samples. It is unclear whether these termites genuinely lack archaea or if this reflects low sequencing depth recovering only abundant community members.

annoSnake functional annotation summarised potential metabolic pathways of interest (Fig. 3). In all samples, annoSnake recovered several genes related to the dissimilatory sulfate reduction pathway: *aprA, aprB*, and *dsrAB*. Sulfate concentration and abundance of sulfate-reducing bacteria are generally low in termite guts (Brune and Okhuma 2011, Arora et al. 2022). Few genes related to methanogenesis, such as *mcrABG*, were found. This is expected, as methanogenesis is largely constrained to anaerobic methanogenic archaea (Thauer et al. 2008, Brune 2010), which were not detected here (Fig. 1).

To target acetogens in the data, we specified genes corresponding to seven enzymes of the Wood-Ljungdahl pathway: *fdhF, fhs, folD, metF, acsABCDE, pta*, and *ack*. Although *fhs* and *folD* were not found in our metagenomes, the presence of *fdhF* and *acsABCDE* is predictive of reductive acetogenesis (Fig. 3), known from other termites (Arora et al. 2022) and millipedes (Nweze et al. 2024). This is supported by annoSnake assignment of 15 and 6 MAGs to clostridia and spirochetes, respectively, taxa which include potential acetogens (Okhuma et al. 2005, Mikaelyan et al. 2016).

annoSnake recovered a large set of CAZymes (Fig. 4). *D. tamminensis* gut metagenomes again suggest dietary flexibility, with numerous and abundant glucoside hydrolysases (GHs) in some colonies but not others. Although *D. tamminensis* harvests foodstuffs from wood to grass (Watson and Perry 1981), the observed pattern of GHs in some colonies was consistent with humus/soil-feeding. We found many GHs in the spirochete-dominated *D. gayi* metagenome (D_gayi_BEC329; Fig. 2 and 4), as expected for a leaf-litter harvesting species, and low numbers of GHs in clostridia-dominated *D. gayi* colonies.

Metagenomic binning of the 31 termite gut metagenomes yielded 30 MAGs in total with low (>30% completion and <10% contamination) to high quality (>90% completion and <5% contamination) (Fig. 5), and average relative abundance of 3%. This included 15 MAGs of Bacillota, 1 MAG of Desulfobacterota, 7 MAGs of Fibrobacterota, 1 MAG of Pseudomonadota, and 6 MAGs of Spirochaetota. Interestingly, MAG BEC335_bin.2 from *A. obeuntis* was assigned to genus *Azovibrio* (Pseudomonadota), a diazotrophic endophyte of grasses (Reinhold-Hurek and Hurek 2006). *A. obeuntis* feeds mainly on dead tree roots (Abensperg-Traun 1993), suggesting *Azovibrio* bacteria may be transient, entering the gut as termites moved through soil accessing their woody diet.

**Figure 5.**
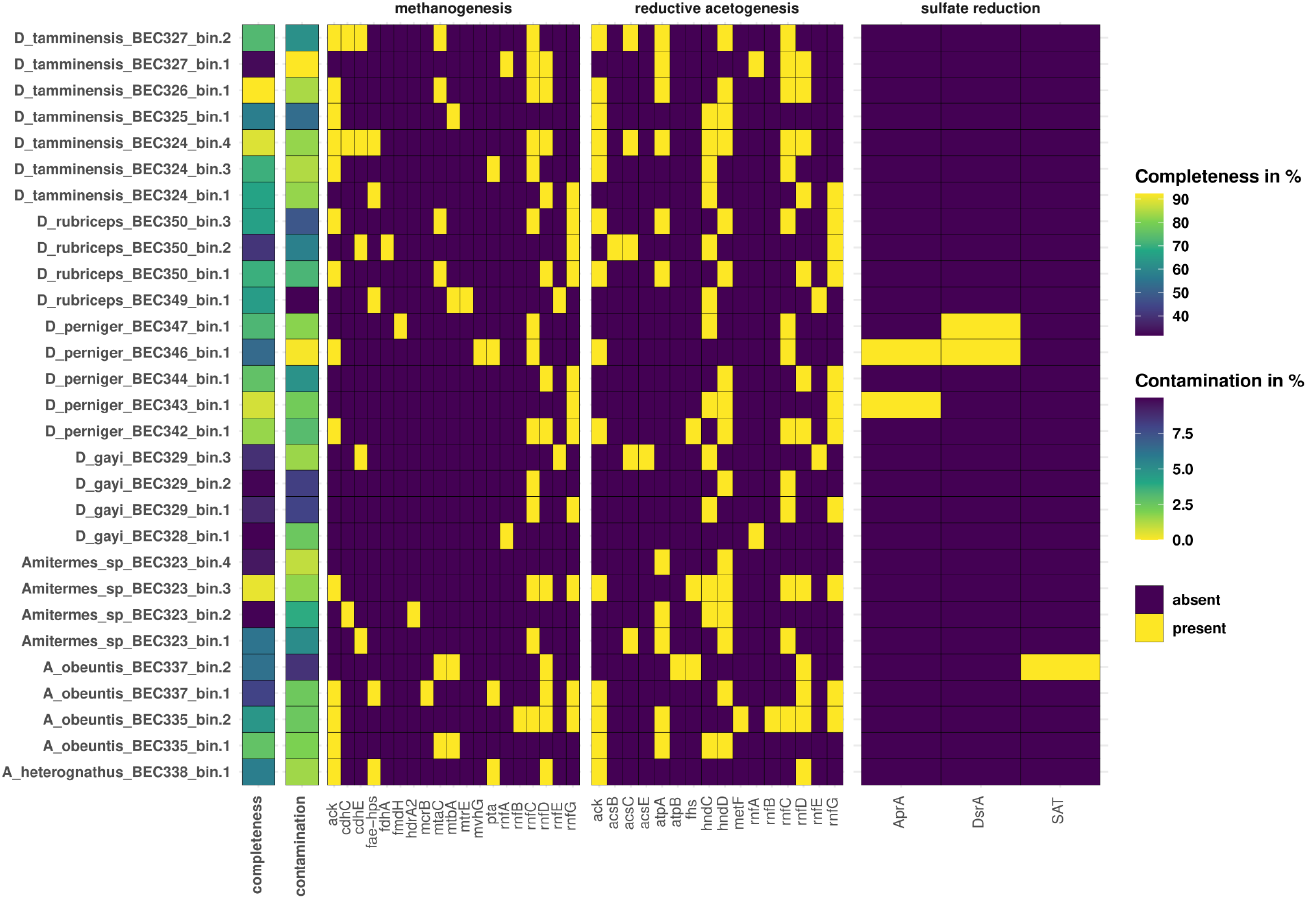
annoSnake representation of lignocellulose digestion metabolic pathway gene presence/absence in MAGs (metagenome-assembled genomes) of 31 termite gut metagenomes (main panels); purple squares indicate absence of genes and yellow squares indicate presence of genes. Leftmost columns: checkM completeness and contamination scores; lighter colours indicate higher completeness and less contamination, respectively.

Altogether, our results show that annoSnake is a powerful tool to facilitate metagenomic analysis. It is easy to implement and highly flexible, allowing the workflow to be easily adapted to different research questions and increasing efficiency in generating novel biological insights from metagenomic sequence data.

## Acknowledgements

The authors hereby declare no conflict of interest. We acknowledge the traditional custodians of Country throughout Australia and pay our respects to Elders past and present. Samples were collected with permission of relevant state authorities.

## References

Abensperg-Traun, M. (1993). A comparison of two methods for sampling assemblages of subterranean, wood-eating termites (Isoptera). Austral Ecology, 18(3), 317–324. 10.1111/j.1442-9993.1993.tb00459.x

Arora, J., Kinjo, Y., Šobotník, J., Buček, A., Clitheroe, C., Stiblik, P., Roisin, Y., Žifčáková, L., Park, Y. C., Kim, K. Y., Sillam-Dussès, D., Hervé, V., Lo, N., Tokuda, G., Brune, A., & Bourguignon, T. (2022). The functional evolution of termite gut microbiota. Microbiome, 10(1), 78. 10.1186/s40168-022-01258-3

Brune, A. (2010). Methanogenesis in the Digestive Tracts of Insects. In K. N. Timmis (Ed.), Handbook of Hydrocarbon and Lipid Microbiology (pp. 707–728). Springer. 10.1007/978-3-540-77587-4_56

Brune, A., & Ohkuma, M. (2011). Role of the Termite Gut Microbiota in Symbiotic Digestion. In D. E. Bignell, Y. Roisin, & N. Lo (Eds.), Biology of Termites: A Modern Synthesis (pp. 439–475). Springer Netherlands. 10.1007/978-90-481-3977-4_16

Buchfink, B., Reuter, K., & Drost, H.-G. (2021). Sensitive protein alignments at tree-of-life scale using DIAMOND. Nature Methods, 18(4), 366–368. 10.1038/s41592-021-01101-x

Calle, M. L. (2019). Statistical Analysis of Metagenomics Data. Genomics & Informatics, 17(1), e6. 10.5808/GI.2019.17.1.e6

Chaumeil, P.-A., Mussig, A. J., Hugenholtz, P., & Parks, D. H. (2022). GTDB-Tk v2: Memory friendly classification with the genome taxonomy database. Bioinformatics, 38(23), 5315–5316. 10.1093/bioinformatics/btac672

Coventry, R. J., Holt, J. A., & Sinclair, D. F. (1988). Nutrient cycling by mound building termites in low fertility soils of semi-arid tropical Australia. Soil Research, 26(2), 375–390. 10.1071/sr9880375

Dong, X., & Strous, M. (2019). An Integrated Pipeline for Annotation and Visualization of Metagenomic Contigs. Frontiers in Genetics, 10. 10.3389/fgene.2019.00999

Evans, T. A., Dawes, T. Z., Ward, P. R., & Lo, N. (2011). Ants and termites increase crop yield in a dry climate. Nature Communications, 2(1), 262. 10.1038/ncomms1257

Ferrar, P., & Watson, J. A. L. (1970). Termites (Isoptera) Associated with Dung in Australia. Australian Journal of Entomology, 9(2), 100–102. 10.1111/j.1440-6055.1970.tb00778.x

Gay, F. J. (1968). A contribution to the systematics of the genus Amitermes (Isoptera: Termitidae) in Australia. Australian Journal of Zoology, 16(3), 405–457. 10.1071/zo9680405

Kanehisa, M., Furumichi, M., Sato, Y., Kawashima, M., & Ishiguro-Watanabe, M. (2023). KEGG for taxonomy-based analysis of pathways and genomes. Nucleic Acids Research, 51(D1), D587–D592. 10.1093/nar/gkac963

Kanehisa, M., & Goto, S. (2000). KEGG: Kyoto encyclopedia of genes and genomes. Nucleic Acids Research, 28(1), 27–30. 10.1093/nar/28.1.27

Kang, D. D., Li, F., Kirton, E., Thomas, A., Egan, R., An, H., & Wang, Z. (2019). MetaBAT 2: An adaptive binning algorithm for robust and efficient genome reconstruction from metagenome assemblies. PeerJ, 7, e7359. 10.7717/peerj.7359

Krishna, K., Grimaldi, D. A., Krishna, V., & Engel, M. S. (2013). Treatise on the Isoptera of the world. (Bulletin of the American Museum of Natural History, no. 377). https://digitallibrary.amnh.org/handle/2246/6430

Kultima, J. R., Coelho, L. P., Forslund, K., Huerta-Cepas, J., Li, S. S., Driessen, M., Voigt, A. Y., Zeller, G., Sunagawa, S., & Bork, P. (2016). MOCAT2: A metagenomic assembly, annotation and profiling framework. Bioinformatics, 32(16), 2520–2523. 10.1093/bioinformatics/btw183

Lema, N. K., Gemeda, M. T., & Woldesemayat, A. A. (2023). Recent Advances in Metagenomic Approaches, Applications, and Challenges. Current Microbiology, 80(11), 347. 10.1007/s00284-023-03451-5

Li, D., Liu, C.-M., Luo, R., Sadakane, K., & Lam, T.-W. (2015). MEGAHIT: An ultra-fast single-node solution for large and complex metagenomics assembly via succinct de Bruijn graph. Bioinformatics, 31(10), 1674–1676. 10.1093/bioinformatics/btv033

Mallawaarachchi, V., & Lin, Y. (2022). Accurate Binning of Metagenomic Contigs Using Composition, Coverage, and Assembly Graphs. Journal of Computational Biology, 29(12), 1357–1376. 10.1089/cmb.2022.0262

Mikheenko, A., Saveliev, V., & Gurevich, A. (2016). MetaQUAST: Evaluation of metagenome assemblies. Bioinformatics, 32(7), 1088–1090. 10.1093/bioinformatics/btv697

Mistry, J., Chuguransky, S., Williams, L., Qureshi, M., Salazar, G. A., Sonnhammer, E. L. L., Tosatto, S. C. E., Paladin, L., Raj, S., Richardson, L. J., Finn, R. D., & Bateman, A. (2021). Pfam: The protein families database in 2021. Nucleic Acids Research, 49(D1), D412–D419. 10.1093/nar/gkaa913

Mölder, F., Jablonski, K. P., Letcher, B., Hall, M. B., Tomkins-Tinch, C. H., Sochat, V., Forster, J., Lee, S., Twardziok, S. O., Kanitz, A., Wilm, A., Holtgrewe, M., Rahmann, S., Nahnsen, S., & Köster, J. (2021). Sustainable data analysis with Snakemake (10:33). F1000Research. 10.12688/f1000research.29032.2

Nam, N. N., Do, H. D. K., Loan Trinh, K. T., & Lee, N. Y. (2023). Metagenomics: An Effective Approach for Exploring Microbial Diversity and Functions. Foods, 12(11), 10.3390/foods12112140

New, F. N., & Brito, I. L. (2020). What Is Metagenomics Teaching Us, and What Is Missed? Annual Review of Microbiology, 74, 117–135. 10.1146/annurev-micro-012520-072314

Noble, J. C., Müller, W. J., Whitford, W. G., & Pfitzner, G. H. (2009). The significance of termites as decomposers in contrasting grassland communities of semi-arid eastern Australia. Journal of Arid Environments, 73(1), 113–119. 10.1016/j.jaridenv.2008.08.004

Nweze, J. E., Šustr, V., Brune, A., & Angel, R. (2024). Functional similarity, despite taxonomical divergence in the millipede gut microbiota, points to a common trophic strategy. Microbiome, 12(1), 16. 10.1186/s40168-023-01731-7

Parks, D. H., Chuvochina, M., Chaumeil, P.-A., Rinke, C., Mussig, A. J., & Hugenholtz, P. (2020). A complete domain-to-species taxonomy for Bacteria and Archaea. Nature Biotechnology, 38(9), 1079–1086. 10.1038/s41587-020-0501-8

Parks, D. H., Chuvochina, M., Rinke, C., Mussig, A. J., Chaumeil, P.-A., & Hugenholtz, P. (2022). GTDB: An ongoing census of bacterial and archaeal diversity through a phylogenetically consistent, rank normalized and complete genome-based taxonomy. Nucleic Acids Research, 50(D1), D785–D794. 10.1093/nar/gkab776

Parks, D. H., Imelfort, M., Skennerton, C. T., Hugenholtz, P., & Tyson, G. W. (2015). CheckM: Assessing the quality of microbial genomes recovered from isolates, single cells, and metagenomes. Genome Research, 25(7), 1043–1055. 10.1101/gr.186072.114

Patro, R., Duggal, G., Love, M. I., Irizarry, R. A., & Kingsford, C. (2017). Salmon provides fast and bias-aware quantification of transcript expression. Nature Methods, 14(4), 417–419. 10.1038/nmeth.4197

Platell, G. A. M. M. (2019). Exploring intraspecific variation in the gut communities of Western Australian endemic termites (lsoptera, Termitidae) as a foundation for future local biofuel initiatives [Doctoral Thesis].

Quinn, T. P., Richardson, M. F., Lovell, D., & Crowley, T. M. (2017). propr: An R-package for Identifying Proportionally Abundant Features Using Compositional Data Analysis. Scientific Reports, 7(1), 16252. 10.1038/s41598-017-16520-0

Ruiz-Perez, C. A., Conrad, R. E., & Konstantinidis, K. T. (2021). MicrobeAnnotator: A user-friendly, comprehensive functional annotation pipeline for microbial genomes. BMC Bioinformatics, 22(1), 11. 10.1186/s12859-020-03940-5

Seemann, T. (2014). Prokka: Rapid prokaryotic genome annotation. Bioinformatics, 30(14), 2068–2069. 10.1093/bioinformatics/btu153

Setubal, J. C. (2021). Metagenome-assembled genomes: Concepts, analogies, and challenges. Biophysical Reviews, 13(6), 905–909. 10.1007/s12551-021-00865-y

Sievert, C. (2020). Interactive Web-Based Data Visualization with R, plotly, and shiny. Chapman and Hall/CRC. 10.1201/9780429447273

Søndergaard, D., Pedersen, C. N. S., & Greening, C. (2016). HydDB: A web tool for hydrogenase classification and analysis. Scientific Reports, 6(1), 34212. 10.1038/srep34212

Sunagawa, S., Mende, D. R., Zeller, G., Izquierdo-Carrasco, F., Berger, S. A., Kultima, J. R., Coelho, L. P., Arumugam, M., Tap, J., Nielsen, H. B., Rasmussen, S., Brunak, S., Pedersen, O., Guarner, F., de Vos, W. M., Wang, J., Li, J., Doré, J., Ehrlich, S. D., … Bork, P. (2013). Metagenomic species profiling using universal phylogenetic marker genes. Nature Methods, 10(12), Article 12. 10.1038/nmeth.2693

Thauer, R. K., Kaster, A.-K., Seedorf, H., Buckel, W., & Hedderich, R. (2008). Methanogenic archaea: Ecologically relevant differences in energy conservation. Nature Reviews Microbiology, 6(8), 579–591. 10.1038/nrmicro1931

Uritskiy, G. V., DiRuggiero, J., & Taylor, J. (2018). MetaWRAP—a flexible pipeline for genome-resolved metagenomic data analysis. Microbiome, 6(1), 158. 10.1186/s40168-018-0541-1

Watson, J. A. L., & Perry, D. H. (1981). The Australian harvester termites of the genus Drepanotermes (Isoptera: Termitinae). Australian Journal of Zoology Supplementary Series, 29(78), 1–153. 10.1071/ajzs078

Wickham, H. (2016). ggplot2: Elegant Graphics for Data Analysis. Springer Publishing New York

Wu, Y.-W., Simmons, B. A., & Singer, S. W. (2016). MaxBin 2.0: An automated binning algorithm to recover genomes from multiple metagenomic datasets. Bioinformatics, 32(4), 605–607. 10.1093/bioinformatics/btv638

Xia, Y., & Sun, J. (2021). Statistical Data Analysis of Microbiomes and Metabolomics. American Chemical Society. 10.1021/acsinfocus.7e5035

Yin, Y., Mao, X., Yang, J., Chen, X., Mao, F., & Xu, Y. (2012). dbCAN: A web resource for automated carbohydrate-active enzyme annotation. Nucleic Acids Research, 40(W1), W445–W451. 10.1093/nar/gks479

Yoo, A. B., Jette, M. A., & Grondona, M. (2003). SLURM: Simple Linux Utility for Resource Management. In D. Feitelson, L. Rudolph, & U. Schwiegelshohn (Eds.), Job Scheduling Strategies for Parallel Processing (pp. 44–60). Springer. 10.1007/10968987_3

Zhou, Y., Liu, M., & Yang, J. (2022). Recovering metagenome-assembled genomes from shotgun metagenomic sequencing data: Methods, applications, challenges, and opportunities. Microbiological Research, 260, 127023. 10.1016/j.micres.2022.127023

