## Supporting Information for "annoSnake: a Snakemake workflow for taxonomic and functional annotation of metagenomes and metagenome-assembled genomes (MAGs)"

### **DNA extraction and sequencing**

We isolated guts from three workers from each of 31 colonies representing nine *Amitermes* and *Drepanotermes* species (originally collected for Clement et al. 2021 and Heimburger et al. 2022; Tab. S1)*.* Pooled guts were homogenised using the Precellys Evolution Touch homogenizer (Bertin Instruments, France), then DNA extracted using the Qiagen DNeasy PowerSoil kit (Qiagen, Germany). Samples were prepared for PCR-free DNA sequencing using Illumina’s Nextera XT kit. Paired-end libraries (250 bp) were sequenced on a NextSeq 550 machine (Illumina, USA) using a mid-output sequencing kit at George Washington University’s Sequencing Core. Raw paired-end reads used in the workflow demonstration can be found under https://figshare.com/s/59c0bbaacf2f8e573bf2.

**Supplementary Table 1**	Summary information including sample IDs, species names, feeding habits (reference), GPS coordinates of sampling locations, and number of assembled MAGs.

| **Sample ID** | **Species** | **Feeding habits** | **Latitude** | **Longitude** | **# of MAGs** |
| --- | --- | --- | --- | --- | --- |
| **BEC2** | *A. laurensis* | grass-litter/dung (Lee and Wood 1971) | -16.63 | 145.23 | 0 |
| **BEC5** | *A. laurensis* | grass-litter/dung (Lee and Wood 1971) | -16.7 | 145.26 | 0 |
| **BEC8** | *A. laurensis* | grass-litter/dung (Lee and Wood 1971) | -16.63 | 145.24 | 0 |
| **BEC002** | *A. laurensis* | grass-litter/dung (Lee and Wood 1971) | -16.63 | 145.24 | 0 |
| **BEC323** | *Amitermes* sp | Na | -18.19 | 142.89 | 4 |
| **BEC324** | *D. tamminensis* | wood/grass (Watson and Perry 1981) | -30.96 | 116.61 | 3 |
| **BEC325** | *D.tamminensis* | wood/grass (Watson and Perry 1981) | -30.39 | 116.64 | 1 |
| **BEC326** | *D. tamminensis* | wood/grass (Watson and Perry 1981) | -30.39 | 116.64 | 1 |
| **BEC327** | *D. tamminensis* | wood/grass (Watson and Perry 1981) | -28.49 | 115.64 | 2 |
| **BEC328** | *D. gayi* | leaf-litter (Watson and Perry 1981) | -30.25 | 116.68 | 1 |
| **BEC329** | *D. gayi* | leaf-litter (Watson and Perry 1981) | -32.39 | 117.96 | 3 |
| **BEC330** | *D. gayi* | leaf-litter (Watson and Perry 1981) | -32.39 | 118 | 1 |
| **BEC331** | *A. dentosus* | wood/dung (Ferrar and Watson 1970) | -28.49 | 115.64 | 0 |
| **BEC332** | *A. dentosus* | wood/dung (Ferrar and Watson 1970) | -27.89 | 117.86 | 0 |
| **BEC333** | *A. dentosus* | wood/dung (Ferrar and Watson 1970) | -30.72 | 115.97 | 0 |
| **BEC334** | *Amitermes* sp | Na | -29.54 | 115.64 | 0 |
| **BEC335** | *A. obeuntis* | roots (Abensperg-Traun 1993) | -31.71 | 116.54 | 2 |
| **BEC336** | *A. obeuntis* | roots (Abensperg-Traun 1993) | -32.17 | 116.19 | 0 |
| **BEC337** | *A. obeuntis* | roots (Abensperg-Traun 1993) | -32.44 | 115.97 | 2 |
| **BEC338** | *A. heterognathus* | wood/dung (Ferrar and Watson 1970) | -31.79 | 116.43 | 1 |
| **BEC340** | *D. rubriceps* | grass (Watson and Perry 1981) | -31.31 | 119.66 | 0 |
| **BEC341** | *D. perniger* | grass (Watson and Perry 1981) | -29.28 | 117.7 | 0 |
| **BEC342** | *D. perniger* | grass (Watson and Perry 1981) | -28.1 | 117.84 | 1 |
| **BEC343** | *D. perniger* | grass (Watson and Perry 1981) | -28.15 | 117.7 | 1 |
| **BEC344** | *D. perniger* | grass (Watson and Perry 1981) | -31.27 | 119.65 | 1 |
| **BEC345** | *D. perniger* | grass (Watson and Perry 1981) | -31.27 | 119.65 | 0 |
| **BEC346** | *D. perniger* | grass (Watson and Perry 1981) | -32.5 | 118.26 | 1 |
| **BEC347** | *D. perniger* | grass (Watson and Perry 1981) | -32.26 | 117.92 | 1 |
| **BEC348** | *D. rubriceps* | grass (Watson and Perry 1981) | -16.58 | 144.92 | 0 |
| **BEC349** | *D. rubriceps* | grass (Watson and Perry 1981) | -16.55 | 144.94 | 1 |
| **BEC350** | *D. rubriceps* | grass (Watson and Perry 1981) | -16.61 | 145.23 | 3 |
|  |  |  |  |  | 30 |

**References**

Abensperg-Traun, M. (1993). A comparison of two methods for sampling	assemblages of subterranean, wood-eating termites (Isoptera). Australian	Journal of Ecology, 18(3), 317–324. https://doi.org/10.1111/j.1442-	9993.1993.tb00459.x

Clement, R. A., Flores-Moreno, H., Cernusak, L. A., Cheesman, A. W., Yatsko, A. R.,	Allison, S. D., Eggleton, P., & Zanne, A. E. (2021). Assessing the Australian	Termite Diversity Anomaly: How Habitat and Rainfall Affect Termite	Assemblages. *Frontiers in Ecology and Evolution*, *9*, 237.	<https://doi.org/10.3389/fevo.2021.657444>

Ferrar, P., & Watson, J. A. L. (1970). Termites (isoptera) Associated with Dung in	Australia. Australian Journal of Entomology, 9(2), 100–102.	https://doi.org/10.1111/j.1440-6055.1970.tb00778.x

Heimburger, B., Schardt, L., Brandt, A., Scheu, S., & Hartke, T. R. (2022). Rapid	diversification of the Australian *Amitermes* group during late Cenozoic climate	change. *Ecography*, *2022*(9), e05944. <https://doi.org/10.1111/ecog.05944>

Lee, K. L. & Wood, T. G. (1971) Physical and chemical effects on soils of some	Australian termites and their pedological significance. Pedobiologia, 11(5),	376-409.

Watson, J. A. L., & Perry, D. H. (1981). The Australian harvester termites of the	genus	*Drepanotermes* (Isoptera: Termitinae). Australian Journal of Zoology	Supplementary Series, 29(78), 1–153. https://doi.org/10.1071/ajzs078
